# Structure of the virulence-associated *Neisseria meningitidis* filamentous bacteriophage MDAΦ

**DOI:** 10.1101/2024.10.02.616247

**Authors:** Jan Böhning, Miles Graham, Mathieu Coureuil, Abul K. Tarafder, Julie Meyer, Xavier Nassif, Emmanuelle Bille, Tanmay A. M. Bharat

**Affiliations:** Structural Studies Division, MRC Laboratory of Molecular Biology, Francis Crick Avenue, Cambridge CB2 0QH, United Kingdom; Université Paris Cité, INSERM U1151, CNRS UMR8253, Institut Necker-Enfants Malades, F-75015 Paris, France

## Abstract

*Neisseria meningitidis* is a human commensal bacterium that can opportunistically invade the bloodstream and cross the blood-brain barrier, where it can cause septicaemia and meningitis. These diseases, if left untreated, can be lethal within hours. Hyperinvasive *N. meningitidis* strains often express a genomically encoded filamentous bacteriophage called MDAΦ, which promotes colonisation of mucosal host surfaces to facilitate invasion. How this phage is organised and how it promotes biofilm formation and infection at the molecular level is unclear. Here, we present an electron cryomicroscopy structure of the MDA phage, showing that MDAΦ is a class I filamentous inovirus, with the major capsid protein arranged within the phage as a highly curved and densely packed α-helix. Comparison with other filamentous bacteriophages offers clues about inoviral genome encapsidation mechanisms, providing a framework for understanding the evolutionary diversity of inoviruses. A disordered, N-terminal segment in the major capsid protein presents hydrophobic patches on the surface of assembled phage particles, which, together with electron cryotomography data of phage bundles, furnishes a structural rationale for phage-phage interactions that were seen previously in an epithelium adhesion infection model of *N. meningitidis*. Taken together, our results shed light on the structure, organisation, and higher order assembly of a biomedically relevant phage encoded in the genome of a human pathogen. Molecular insights gleaned from this study increase our understanding of phage evolution, phage-mediated bacterial adhesion and pathogenicity.

## Introduction

*Neisseria meningitidis* is one of the major causes of bacterial meningitis, which can result in neurological damage and death within hours if untreated (1). *N. meningitidis* is found commensally in the throat, but can invade the bloodstream to cause septicaemia and/or invasive meningeal disease after crossing the blood–brain barrier (2). To become septicaemic, bacteria must first colonize the mucosal surfaces of the nasopharynx, after which they can invade the bloodstream (3-5). Hence, the factors promoting colonization of host mucosal surfaces are of particular importance for understanding disease progression (4-7).

Previous studies found that an 8 kilobase genetic island encoding a filamentous bacteriophage, termed MDAΦ (short for <u>m</u>eningococcal <u>d</u>isease-<u>a</u>ssociated bacteriophage), was a common feature in hyperinvasive populations of *N. meningitidis* (8, 9). Filamentous inoviruses such as MDAΦ are a genus of bacteriophages consisting of a single-stranded DNA (ssDNA) genome that is surrounded by an α-helical capsid, resulting in rod-like phage particles up to several micrometres in length but less than ten nanometres in diameter (10). While employing their host’s resources to replicate, filamentous phages are often continuously secreted into the environment without host lysis. In some cases, the phage is symbiotic with its bacterial host and secretion of the phage provides benefits to its host bacterium (11-13). For example, the human pathogen *Pseudomonas aeruginosa* encodes a phage, Pf4, which is secreted into the biofilm matrix to form protective assemblies around cells (14, 15). In *Vibrio cholerae*, the bacteriophage CTXΦ is a vital feature of virulent strains, carrying the cholera toxin genes responsible for key aspects of the etiology of cholera (16).

In *N. meningitidis*, MDAΦ particles, after being secreted into the extracellular milieu, assemble into filament bundles that promote cohesion between cells, thus promoting the formation of biofilms (17). In a previously proposed model, a layer of heavily piliated bacteria initiate adhesion to host epithelial cells, while subsequent layers of bacteria are encased in MDAΦ filaments that promote the formation of a thick biofilm (17). Promoting colonization of the mucosal surfaces in the nasopharynx increases the likelihood of subsequent invasion, suggesting a direct link between pathogenicity and the presence of the MDA bacteriophage (8, 9).

Despite the ubiquity of filamentous bacteriophages (18), and considering the important roles they play in many diseases (8, 12, 16), it is surprising that there is limited atomic structural information on inoviral phages themselves. Based on the symmetrical arrangement of the major capsid protein (MCP), inoviruses have been classified into two types: in class I inoviruses, the MCP is arranged helically with an additional pentameric rotational (C5) symmetry around the helical axis within the phage, while class II inoviruses have a helically arranged MCP with no additional rotational symmetry around the helical axis (10, 19, 20). All high-resolution structural work on inoviral capsids has been performed on a limited set of phages: the class I Pf phages (14), and the class II Ff phages (15, 21, 22), as well as on the Ff-related IKe phage (23). Little is known about the structure of filamentous bacteriophages outside of these families. It remains unknown which class of phage MDAΦ falls under, or what structural features of its capsid make it able to bundle into higher order assemblies that promote biofilm formation during infection.

Here, we present the electron cryomicroscopy (cryo-EM) structure of native MDA inoviral bacteriophages isolated from *N. meningitidis*, revealing significant differences to previously solved inovirus capsid structures. Based on our structural data, together with previously available sequence information, we illuminate the links between capsid structure and genome arrangement, thus allowing us to classify inoviral genomes as circular or linear simply based on MCP sequence. We combine our analysis of the phage structure with electron cryotomography (cryo-ET) of phage bundles, which enables us to understand the rationale of phage-phage interactions, providing a structural basis for comprehending MDA phage-mediated virulence that depends on phage bundling.

## Results

### Cryo-EM structure of the MDAΦ capsid

To determine the structure of the MDAΦ capsid, we natively expressed and isolated phages from *N. meningitidis* (Methods). MDAΦ particles were then concentrated for cryo-EM sample preparation. Images of the concentrated specimen show elongated rod-like particles consistent with filamentous bacteriophages from previous studies (14, 15, 23) (Figure 1A). Two-dimensional class averages show a helical arrangement of the phage (Figure 1A). We estimated the helical parameters from two-dimensional class averages and employed realspace helical refinement (24) to solve the structure of the MDAΦ MCP to 3.7 Å resolution (Figure 1, Figure S1-2, Movie S1). The resulting cryo-EM map reveals α-helical MCPs packed into a tubular arrangement with an overall diameter of ∼62 Å and a central pore with a ∼20 Å diameter (Figure 1, Figure S3). MCPs are arranged with a pentameric (C5) rotational symmetry around the filament axis, with a rise of 13.9 Å and 42.3° rotation per subunit, establishing MDAΦ as a class I inovirus.

**Figure 1:**
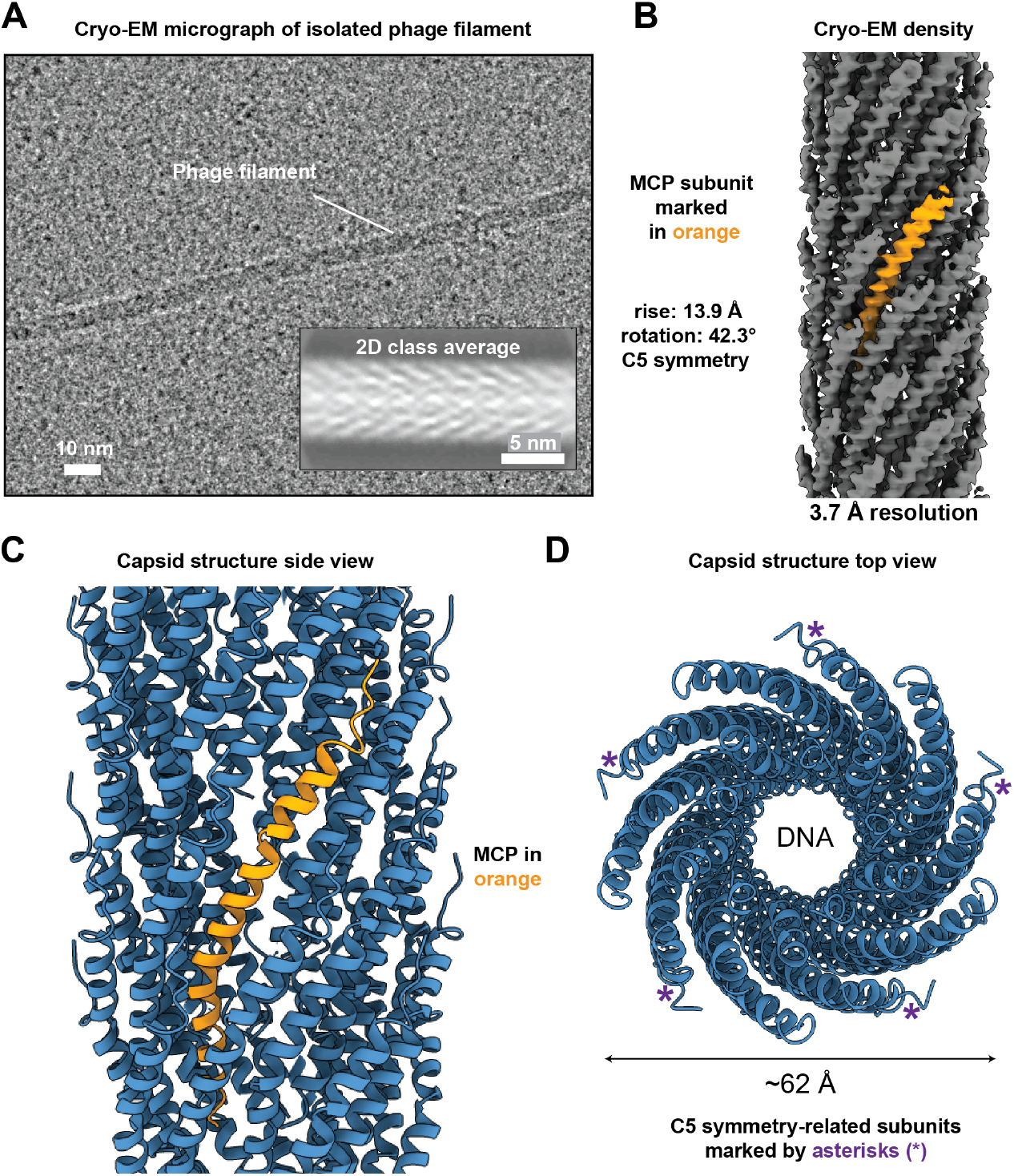
Cryo-EM structure of the filamentous bacteriophage MDAΦ. **A)** Cryo-EM image of MDAΦ particles. Inset: 2D class average of MDAΦ, showing the arrangement of the α-helical capsid protein. **B)** Cryo-EM density map, with a single major capsid protein (MCP) subunit marked. **C-D)** Side and top view of the atomic model of the MDAΦ capsid, with a single MCP subunit marked in orange in panel C.

Each MCP of MDAΦ adopts a bent architecture with significantly higher curvature than has been seen in previous inovirus capsid structures, showing an angle of ∼45° between the N- and the C-terminal segments of the MCP (Figure 2A). The first six residues of the MCP are poorly resolved with smeared density in the cryo-EM map, suggesting flexibility of this region (Figures 1 and S1-S2). This observation is in line with other class I bacteriophages such as Ff phages, where the N-terminal part of the MCP was found to be disordered in previous cryo-EM structures (15, 21, 22). Consistent with past structural data on inoviruses, the MCP of MDAΦ contains numerous hydrophobic residues in the central segment, which form an extensive hydrophobic network through tight packing with other MCP subunits around the filament axis (Figure 2B).

**Figure 2:**
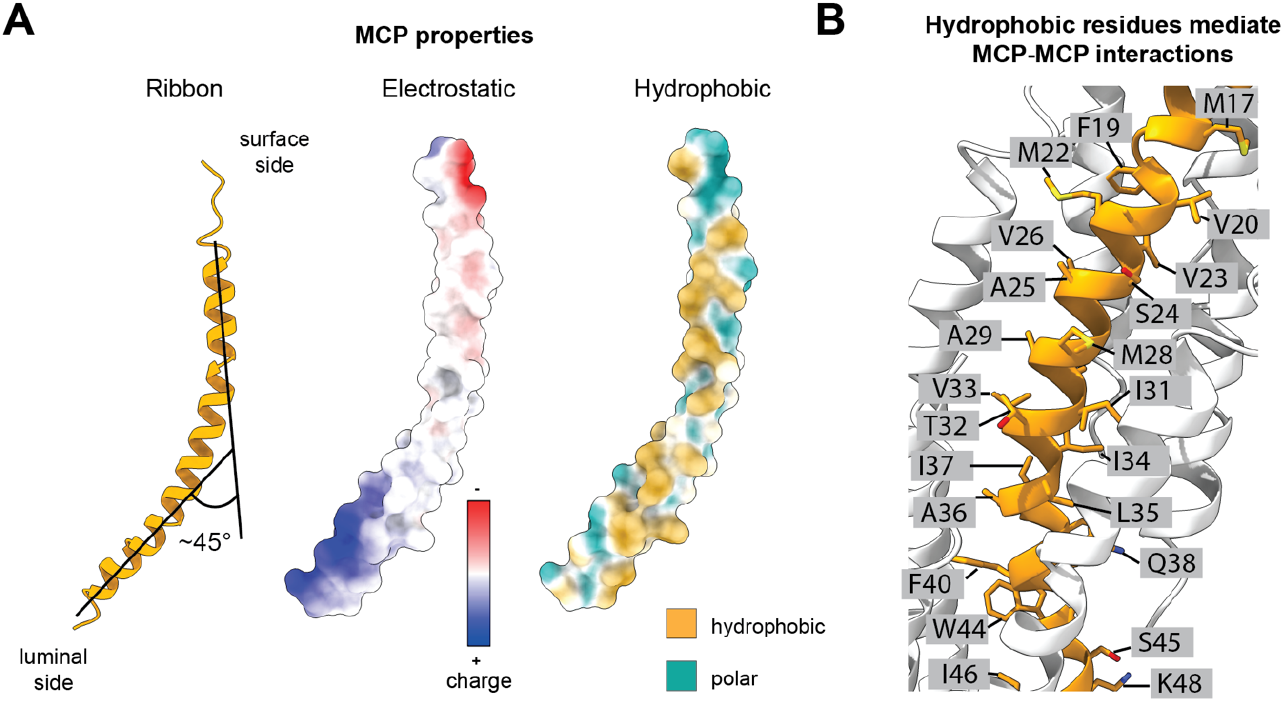
Structural characteristics of the MDAΦ MCP. **A)** Side view of the MCP in ribbon depiction, electrostatic surface depiction, and hydrophobic surface depiction. **B)** Side chains in the central region of the MCP, illustrating that central MCP residues are predominantly hydrophobic, mediating subunit-subunit interactions in the curved surface lattice of the phage.

### Comparison of inovirus structures

The helical parameters of the MDAΦ capsid (rise: 13.9 Å, right-handed rotation 42.3°) are markedly different from other C5-symmetric class I phages such as the Ff group of phages (fd phage rise 16.6 Å, fd phage rotation 36.4°-37.5°) and IKe (rise 16.8 Å, rotation 38.5°), with a lower rise resulting in a comparably higher number of MCP proteins in any given capsid segment. The helical rise of the pentameric, C5-symmetric MDAΦ subunit is also smaller compared to the rise of a comparable, five-MCP subunit of the C1-symmetric class II phage Pf4 (rise of 15.7 Å for Pf4 pseudo-pentameric unit versus 13.9 Å for MDAΦ pentameric unit, Figure 3).

**Figure 3:**
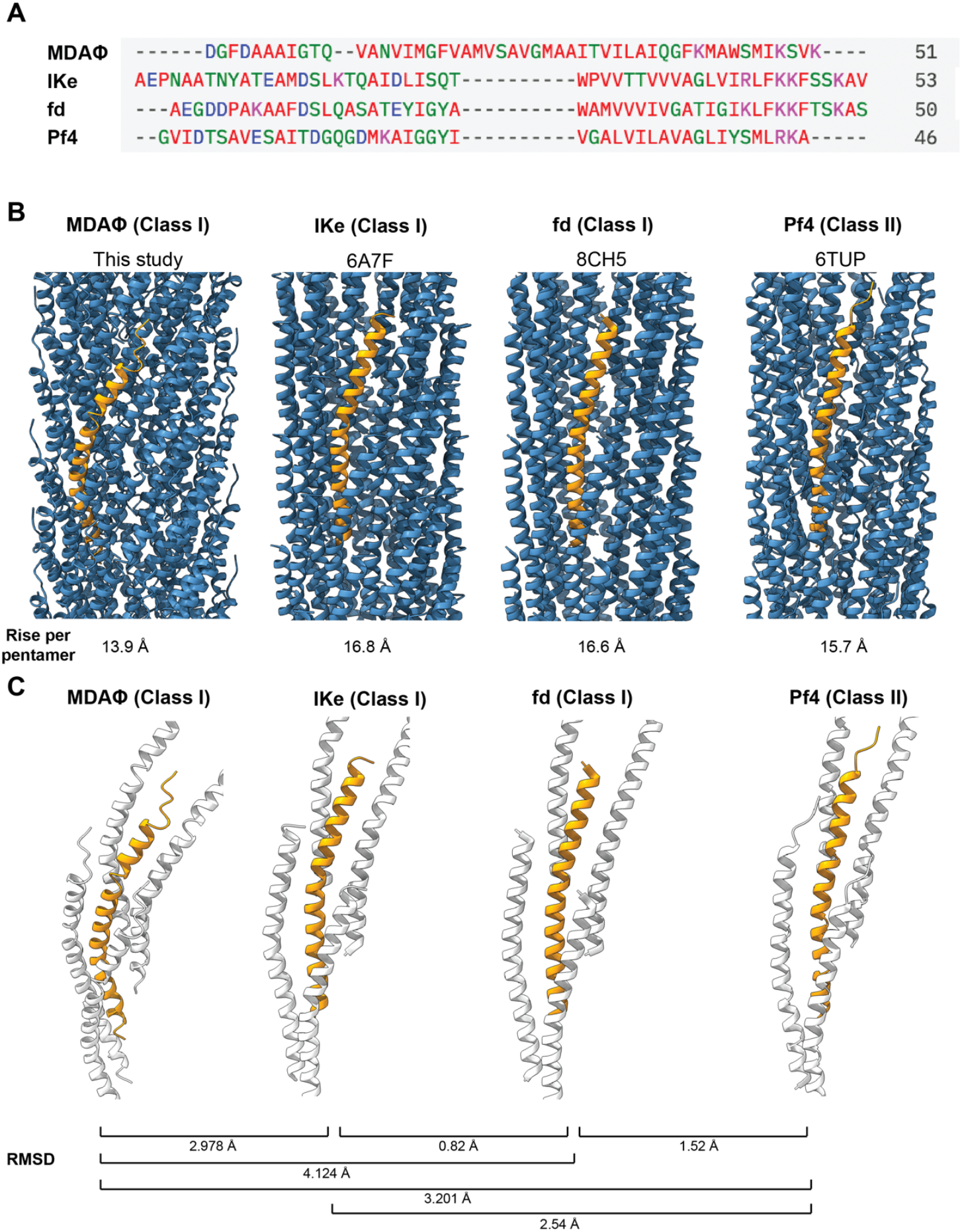
Comparison of the MDAΦ MCP with structurally characterised inoviral MCPs. **A)** Clustal Omega sequence alignment of the mature MCP sequences of MDAΦ, IKe, fd and Pf4. Color code: Blue: acidic residues; violet: basic residues; red: aliphatic residues; green: polar residues. **B)** Side view of the phage MCPs, with a single MCP marked in orange. The rise is respectively given for five MCP subunits, which represents the C5-symmetric subunit in the class I bacteriophages MDAΦ, IKe, and fd. **C)** A single MCP subunit is shown with the four closest neighbouring MCP subunits. Root mean square deviation (RMSD) values are comparing single MCPs.

When we compared previously solved inoviral MCP structures, we found that all inoviral MCPs are broadly similar in overall architecture and arrangement, despite large sequence diversity in different classes of inoviruses (Figure 3). Previously solved inoviral structures show a parallel arrangement of the MCPs along the filament axis which result in largely similar inter-subunit interactions.

The MDAΦ MCP, on the other hand, possesses a notably more curved subunit with a higher tilt relative to the filament axis compared with previously solved phage MCPs (Figures 2 and 3). Compared to other class I phages, which contain a highly ordered α-helical segment terminated by a helix-breaking proline residue (15, 22, 23, 25), flexibility in the MDAΦ MCP continuously increases towards the N-terminus as deduced by local resolution measurements, with the first six residues of the MCP not well resolved (Figure S2). On the whole, these characteristics suggest that MDAΦ represents a structurally divergent filamentous inovirus. Little is known about the other protein components of MDAΦ, including those that form the tip proteins terminating the capsid tube and mediating receptor binding. ORF6 (NMA1797) of MDAΦ has been implicated as an adhesion protein akin to G3P in Ff phages (21). It is noteworthy that, using AlphaFold3 modelling (26), we were able to model a putative complex containing five subunits of ORF6, five subunits of ORF7 (NMA1798), together with the MCP (ORF4; NMA1795) (Figure S4). The predicted complex has a highly similar arrangement to a previously published cryo-EM structure of the f1 phage pointy tip (Figure S4) (21), suggesting that the tip architecture may be similar to other class I phages.

### Binding of MCP to the genome

The C-terminus of the MDAΦ MCP, which is exposed to the phage lumen, contains three lysine residues with side chains extending towards the phage genome (Figure 4 and S3). Such positively charged amino acid residues are commonly found at the C-termini of inoviral MCPs, which compensate the negative charge of the ssDNA genome running along the inside of the phage filament (15, 21-23). Two types of DNA genomes spanning the length of the filamentous phage capsid have been described: circular ssDNA and linear ssDNA. For circular ssDNA genomes, the genome loops back on itself, which requires the capsid to compensate twice as much charge compared to linear ssDNA per unit length of the phage. This, in turn, requires additional positive charges in the C-terminus of each MCP for genome encapsidation.

**Figure 4:**
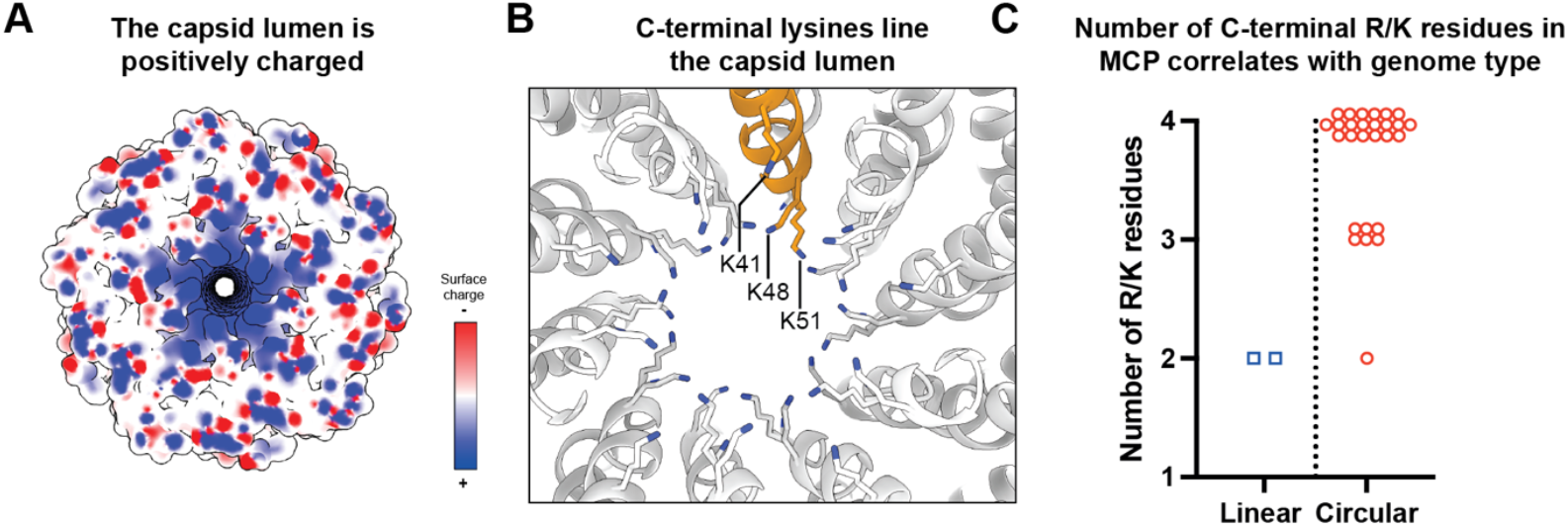
The C-terminus of the MCP coordinates the luminal DNA. **A)** Electrostatic surface depiction of a section of the MDA phage showing positive charge within the phage lumen (blue = positively charged, red = negatively charged). **B)** Three C-terminal lysine residues extend towards the phage genome. **C)** Number of arginine (R) or lysine (K) residues in the C-termini of the MCPs from reference inoviral genomes as annotated by Roux et al. (18).

In previous inoviral MCP structures, either four positively charged residues coordinating a circular ssDNA (Ff and Ike phages) (15, 21-23) or two positively charged residues coordinating a linear ssDNA (Pf4) were observed (14). Density for the ssDNA genome was previously uninterpretable for the class I f1, fd, and IKe phages (15, 21, 23) as well as for the MDA phage in this study (Figure S4), while for the Class II inovirus Pf4, density consistent with a single linear ssDNA was observed (14). We inspected available inoviral reference genomes that were mined previously through exhaustive bioinformatic analyses (18), to assess whether the total number of C-terminal positively charged MCP residues is predictive for the arrangement of the inoviral DNA genome. We only considered reference genomes where the genome type had been previously determined to be linear or circular through direct sequencing of the phage DNA. Indeed, we found that inoviruses with genomes annotated as circular almost exclusively had either 3 or 4 basic residues in the C-terminus of the MCP, while linear inovirus had 2 basic residues in the C-terminus of their MCP (Figure 4). Out of 28 reference phages, there was only a single exception, the inovirus Pf3, the genome of which was annotated to be circular, although its MCP contains only two basic residues. This empirical classification scheme places MDAΦ as a filamentous bacteriophage with a circular ssDNA, which is also consistent with sequencing experiments performed previously (27). This is also consistent with the phage particle length, which, at ∼1200 nm together with the genome size of ∼8 kilobases (27), is consistent with a circular single-stranded genome.

### The MDAΦ MCP exposes hydrophobic residues on the phage surface

Previous studies have suggested that MDAΦ phage filaments bundle together to promote colonization of epithelial cells by promoting cohesion between bacterial cell layers (17). Such filament-bundling-mediated biofilm formation has also been observed in other bacteria (28-30); therefore, understanding filament-filament interactions is important to understand this aggregation process at the cellular level. To probe this further, we imaged bundles of MDA phages present in our sample using cryo-ET, which showed a dense arrangement of phages stacked along their long axes (Figures 5A and S5, Movie S2). A Fourier analysis of the spacing of phages in our data reveals a weak equatorial repeat, orthogonal to the long axis of the bundle, which is close to the diameter of individual phage particles. This repeat agrees with the packing of the phages observed inside the bundles (Figure 5A, box), with longitudinal alignment of the phages. These observations show that tight phage-phage interactions are present in the bundle, but variations in phage-phage spacings are possible.

**Figure 5:**
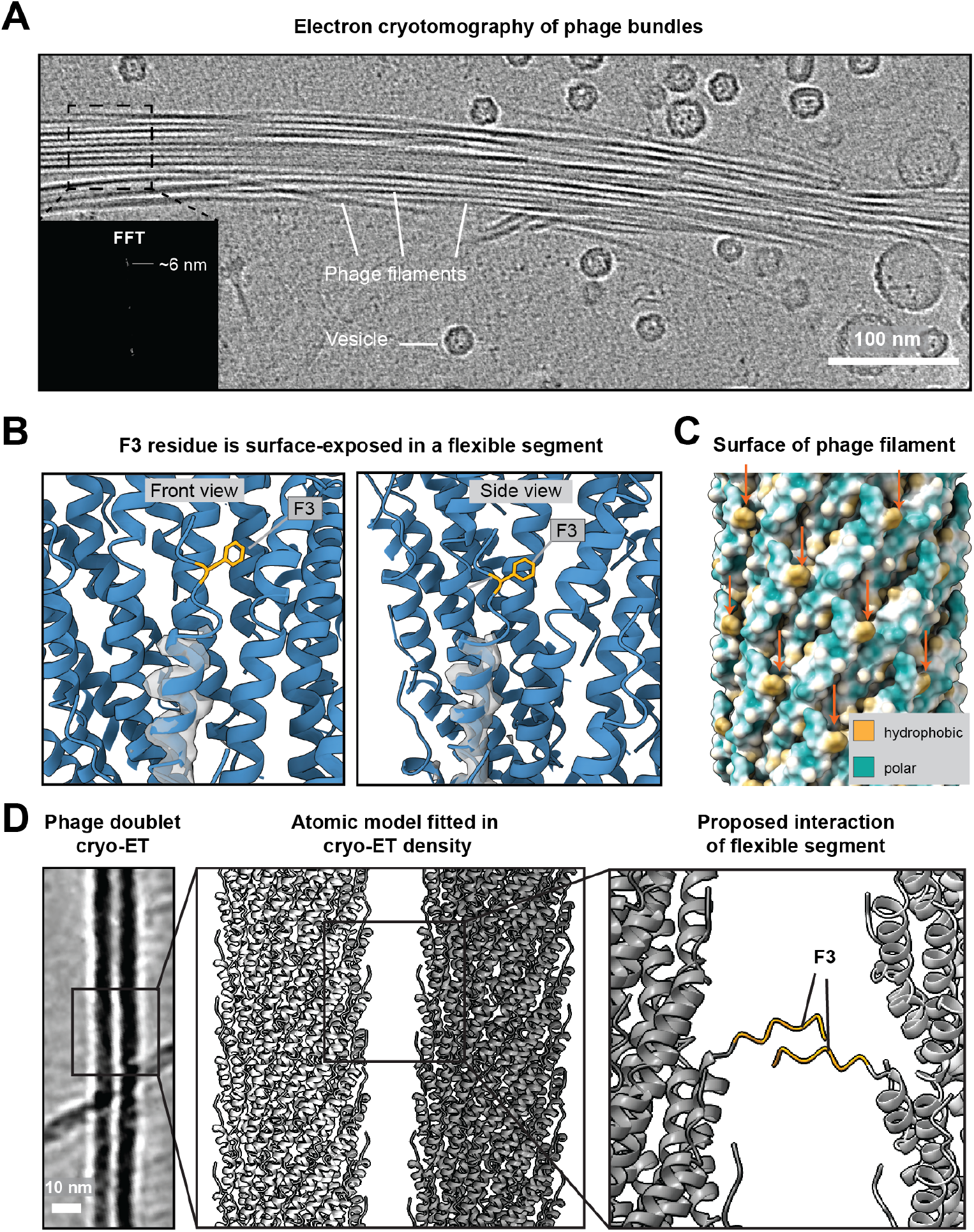
Structural analysis of phage-phage interactions. **A)** Cryo-ET slice depicting a bundle of MDA phages. Inset: Fast Fourier transform (FFT) of a segment of the bundle depicting a range of phage-phage interaction signal around ∼6 nm. The section used to produce the FFT is shown as a dashed box. **B)** An N-terminal segment of the MCP containing the F3 residue (marked in orange) is surface-exposed, and part of the flexible N-terminal segment. The cryo-EM density (grey, transparent) shows that the N-terminal segment containing the F3 residue is poorly resolved and likely flexible. **C)** Hydrophobic surface depiction of the capsid (orange arrows indicate hydrophobic patches caused by F3). **D)** Left: Cryo-ET slice of two phages interacting with each other laterally (‘phage doublet’). Middle: Fit of the atomic model of the MDA phage capsid into the tomographic density. E) Modelling of the flexible N-terminal residues (residues 1-6; orange) reveals putative interactions between N-terminal segments of neighbouring phages. Since bundling occurs spontaneously in MDAΦ, without crowding agents, this provides a structural rationale for phage bundling.

To ascertain how bundling is mediated at the molecular level, we inspected our atomic model of the MDAΦ MCP and noticed that the N-terminal residues 1-6 form a surface-exposed segment with a prominent hydrophobic phenylalanine residue (F3) (Figure 5B) in the poorly resolved, flexible N-terminus of the protein, which is followed by further hydrophobic residues 5-9 (sequence AAAI). Since this short segment containing the F3 residues does not appear to be rigidly constrained by the α-helicity of the MCP, we reasoned it may extend outside the phage into the external medium, where it could mediate binding to other phages to support bundle formation due to avidity effects. Due to the repetitive nature of the MCP (C5 symmetry; rise 13.9 Å, rotation per subunit 42.3°), the F3 residue results in the formation of repetitive hydrophobic patches on the phage outer surface (Figure 5B-C) that are ideally positioned to promote phage-phage interactions, i.e. promote bundling of phages with each other. To probe this hypothesis, we fitted our atomic model into the cryo-ET density of two phages interacting with each other in a doublet (Figure 5D). Our analysis shows that while the phages within the doublet density are spaced too far to interact through their ordered capsids, they are sufficiently close for their flexible N-terminal segments to interact (Figure 5D). While these interactions between the flexible segments cannot be trivially resolved through structural methods, they provide a plausible rationale for the presence of the highly hydrophobic F3 residue in this solvent-exposed flexible segment, not seen in other structurally characterized inoviruses (Figure 3A). The high level of symmetry of the MDAΦ capsid moreover means that interaction through this segment results in high avidity, as the MCP regularly repeats at the phage-phage interface, hinting at a molecular model for phage-phage bundling.

## Discussion

Why most *N. meningitidis* populations remain commensal while others become invasive has been the subject of intensive research (1, 6). A common factor in invasive populations is the expression of MDAΦ genes, resulting in phage production (8, 9), facilitating survival on the nasopharynx epithelium (17), which is a prerequisite for invasion. Our structural data on the biomedically relevant MDA phage shows that MDAΦ is a class I, C5-symmetric bacteriophage with dense MCP packing. Although only a handful in number, previous high-resolution cryo-EM structures of inoviral phages allow us to perform a detailed comparison between various phages. The MCP of MDAΦ adopts an unusually curved arrangement compared to the MCPs of other structurally described class I and II inoviral phages (Figure 3). Interestingly, our comparisons show that fd, another class I bacteriophage, is architecturally closer to the class II bacteriophage Pf4 than to MDAΦ, suggesting that this traditional classification based on the symmetry estimated in fibre diffraction experiments is not fully indicative of structural similarity of phage MCPs. Moreover, given that only model filamentous bacteriophage structures have been described previously, our structure also suggests that a significant structural diversity of filamentous phages likely exists that is yet to be uncovered, previously hinted to by bioinformatic studies (18).

In this study, we found a correlation between the number of positively charged residues at the C-terminus and the genome arrangement of the phage. Our analysis shows that almost all bacteriophages with circular ssDNA have MCPs with 3 or 4 C-terminal lysine or arginine residues, and phages with linear ssDNA – although less common – have 2 basic residues at the C-terminus of the MCP. This fairly simple scheme allows classification of hitherto unknown phages into respective categories without the need for detailed molecular analyses or high-resolution structure determination. The only exception to our proposed scheme was the phage Pf3, which according to this criterion would be predicted to have a linear ssDNA genome. The Pf3 MCP is similar at the sequence level to the Pf4 MCP, which was confirmed via structural methods to have a linear ssDNA genome (14). It is hard to resolve these two observations in this single exception to our proposed classification scheme without further experimentation on the Pf3 phage.

Each MDAΦ MCP subunit contains an unstructured N-terminal segment that presents a hydrophobic phenylalanine residue on the phage surface. While many class I phages have disordered N-terminal tails, highly hydrophobic disordered residues are not present on the surface of other class I bacteriophages (15, 21-23). Due to the highly repetitive nature of the MCP on the phage surface, a high-avidity interaction between neighboring phages could be produced by stacking of these N-terminal phenylalanine residues, suggesting a biochemical mechanism for phage bundling without crowding agents. Mutational analysis of the phage capsid in adhesion models could probe this hypothesis in future studies, although this would be technically complex, requiring mutations in the MDA prophage sequence in the *N. meningitidis* genome, which may affect other aspects of the viral replication cycle including assembly. A further interesting question to be probed is whether the bundles formed by MDAΦ are in direct contact with other biofilm components during infection, which appears plausible but has not been conclusively shown.

Notably, there is precedence for phage bundling providing a beneficial effect to bacterial survival in other species. In *P. aeruginosa* for example, the phage Pf4 is expressed during infections caused by biofilms (31). Pf4 forms bundles around the bacterial cells that prevent diffusion of bactericidal molecules to the bacterial cell surface, promoting antibiotic tolerance (14, 15). Since there are no N-terminal, disordered, hydrophobic residues in the Pf4 MCP, bundling requires a crowding agent such as alginate or hyaluronan, through an entropic effect termed depletion attraction (11). It is conceivable that physiologically relevant crowding agents such as hyaluronan could further enhance MDAΦ phage-phage interactions in a biomedical setting to support the formation of large phage bundles, although this will require experimental validation in future studies. Another beneficial effect of Pf4 phage expression at the sites of *P. aeruginosa* infection is the triggering of human antiviral immune responses, downregulating the antibacterial response, promoting bacterial survival (12, 31). It has been previously shown that MDAΦ bundling reinforces bacterial adhesion (17); whether this also triggers an antiviral response during *Neisseria* infection, thus aiding bacterial survival, remains to be determined. Another notable example of a filamentous phage bolstering virulence is provided by CTXΦ in *V. cholerae*, where the prophage CTX encodes the cholera toxin genes that cause the symptoms of the disease (16). The fact that inoviruses are being employed by numerous bacterial pathogens, all of which appear to exploit the properties of the phage in differing ways, suggests that symbiotic relationships between inoviruses and bacteria may be common. This use of symbiotic phages by bacteria to enhance survival is an area of biomedical research that needs urgent attention in future studies.

## Acknowledgements

This work was supported by the Medical Research Council, as part of United Kingdom Research and Innovation (also known as UK Research and Innovation) [Programme MC_UP_1201/31 to T.A.M.B.]. T.A.M.B. would like to thank the European Molecular Biology Organization, the Wellcome Trust (Grant 225317/Z/22/Z), the Leverhulme Trust, and the Lister Institute for Preventative Medicine for support. We thank Garib Murshudov and Keitaro Yamashita for helpful advice on the model building and Olivia Smith for critical reading of the manuscript. We acknowledge the MRC LMB electron microscopy facility for help with sample preparation and data collection.

## Author contributions

J.B., A.T., X.N., E.B. and T.A.M.B. designed research. J.B., M.G., M.C., A.T.,

J.M., and T.A.M.B. performed research. J.B., M.G., A.T. and T.A.M.B analysed data. J.B., E.B. and T.A.M.B. wrote the manuscript with support from all authors.

## Competing interest statement

The authors declare no competing interests.

## Materials and Methods

### Isolation of phage particles

For the phage isolation, the strain Z5463, formerly designated C396, isolated from the throat of a patient with meningitis in The Gambia in 1983 (32) was used. As previously described (27), bacteria were pelleted from 400 mL of an overnight culture in BHI liquid medium (Condalab) at 37 °C with 5% CO_2_ and agitation. After filtration at 0.45 μm, the supernatant was treated for 2 hours at 20 °C with DNase I (25 μg/mL). Phage was precipitated by addition of 10% NaCl and 20% polyethylene glycol (PEG)6000, and overnight incubation at 4 °C. The phage was then pelleted by centrifugation at 10,000 *g* for 30 minutes and resuspended in PBS (phosphate buffered saline) 1X (with CaCl_2_ and MgCl_2_; Gibco). To further concentrate the phage preparation for cryo-EM, the sample was adjusted to 0.5 M NaCl and 10% (w/v) PEG 6000 before overnight incubation at 4°C. Phage particles were pelleted by centrifugation at 15,000 *g* for 30 minutes. The resulting pellet containing phage was resuspended in PBS followed by dialysis against PBS overnight at 4 °C using 10 kDa MWCO snakeskin dialysis membranes (ThermoFisher).

### Cryo-EM grid preparation

For cryo-EM and cryo-ET grid preparation, previously described protocols were utilized (33). Briefly, 2.5 μL of the concentrated phage was pipetted onto glow discharged Quantifoil grids (Au R1.2/1.3, 400 mesh) and plunge frozen into liquid ethane using a Vitrobot Mark IV (ThermoFisher), with the chamber maintained at 100% relative humidity and 10 °C.

### Cryo-EM image acquisition and processing

Cryo-EM data was acquired in the EER format on a Titan Krios G2 microscope equipped with a Falcon 4 detector, at a nominal magnification of 96,000x using a combined dose of 41 e/Å^*2*^ per movie. For image processing, EER movies were converted to TIF and motion-corrected using the RELION4 implementation of MotionCor2 (34, 35). CTF refinement was performed using CTFFIND4 (36). Phage particles were picked using Topaz (37). Symmetry was determined based on 2D class averages. The correct symmetry resulted in significantly improved resolution and metrics and resulted in resolution of α-helical densities. All classifications and refinements were performed in RELION4 (35, 38). A workflow for the performed image processing steps is shown in Figure S1.

### Atomic model building

An atomic model of the MDA phage capsid was built with Coot using a structural prediction by AlphaFold2 as a starting model. Multiple chains were built to account for subunit-subunit interactions. Refinement was initially performed in PHENIX (39) and then in Servalcat (40) using Servalcat’s helical refinement pipeline. The final refinement was performed in Servalcat. Map and model statistics are shown in Table S1.

### Cryo-ET imaging

Cryo-ET data was acquired on a Titan Krios G4 microscope equipped with a Falcon 4i detector and SelectrisX energy filter, at a nominal magnification at 53,000x and pixel size of 2.39 Å. Tilt series were acquired from -60° to +60° with 3° increments using a dose-symmetric tilt scheme in SerialEM (41) and a total dose of 120 e-/Å^*2*^ over the tilt series at -5 to -7 μm nominal defoci. Tomograms were reconstructed using the etomo package implemented in IMOD (42) using patch tracking and SIRT, or using SART as implemented in AreTomo (43). The Cryo-CARE software package was used for denoising of tomograms (44).

### Phage MCP sequence alignment and genome analysis

Sequence alignment of MCPs was performed using Clustal Omega as implemented by the EBI web server (45). For the analysis of basic C-terminal residues, amino acid sequences of the MCPs in reference genomes as reported by Roux et al. (18) were used, and the occurrence of lysine or arginine residues in the C-terminal 15 residues was assessed. A comprehensive list of analysed genomes and MCPs is given in Table S2.

### Data visualisation

Atomic models were visualised in ChimeraX. FSC plots were created in MATLAB (R2022b), and the genome type plot was created GraphPad Prism and subsequently manually adjusted. Micrographs were filtered in Fiji (46) and displayed using Fiji or IMOD (42). Structural predictions were performed using the AlphaFold3 web server (26). Cryo-ET data was visualised using IMOD.

## Supplementary Figures

**Figure S1:**
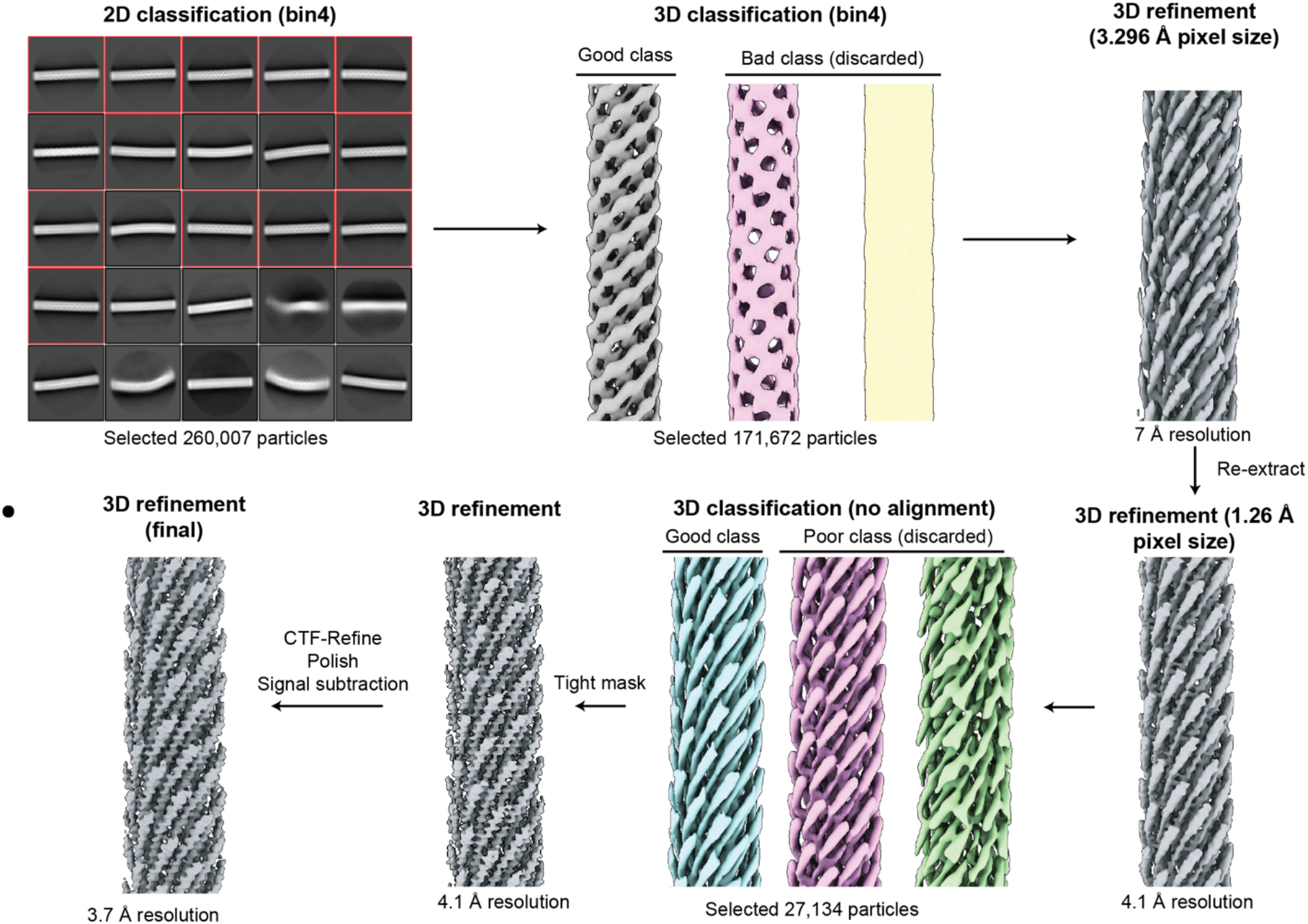
Cryo-EM data processing workflow. Shown are the major classes selected, including particle numbers and further processing steps. Red squares show selected 2D classes; classes not marked were discarded.

**Figure S2:**
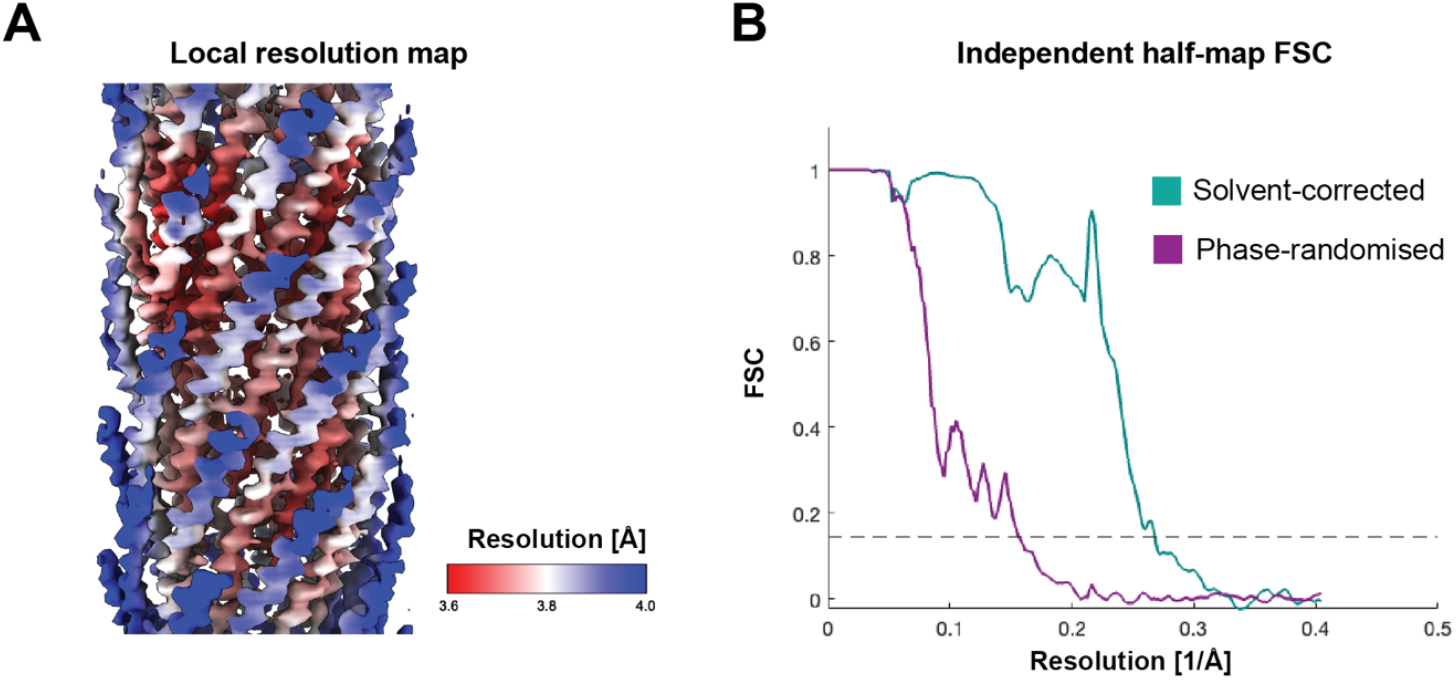
Cryo-EM resolution estimation. **A)** Cryo-EM map coloured by local resolution, scale bar shows resolution values. **B)** Resolution estimation by Fourier shell correlation (FSC) of independently aligned and averaged half-maps, dashed line indicates 0.143 criterion.

**Figure S3:**
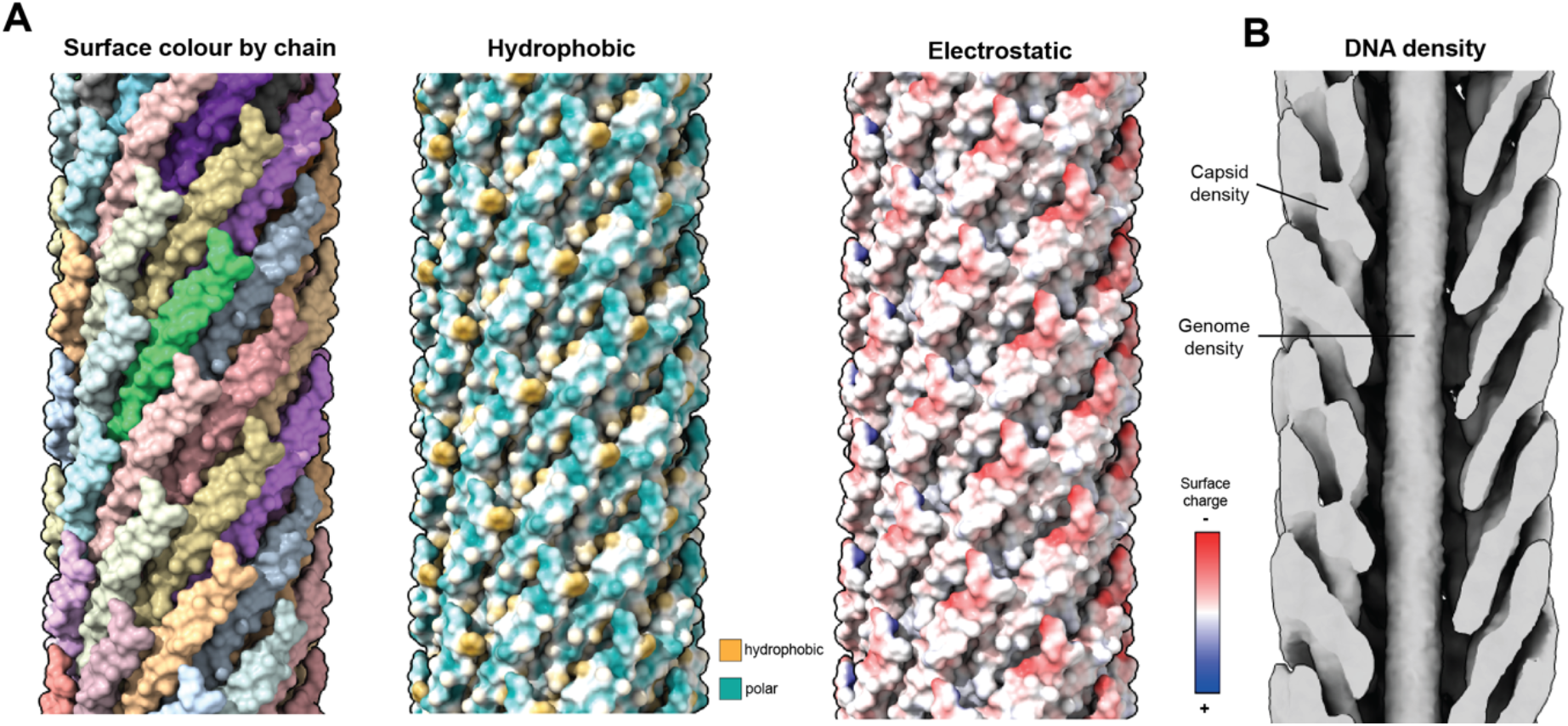
Surface properties and genome density of the MDAΦ map. **A)** Model of the MDAΦ capsid is shown as a surface coloured by chain, as hydrophobic surface depiction (cyan = hydrophilic, yellow = hydrophobic) and electrostatic surface depiction (red = negatively charged, blue = positively charged). **B)** DNA density in the lumen of the MDAΦ phage, in a cryo-EM map produced with C1 symmetry applied, indicating that DNA features are not smeared by the C5 symmetry, because the genome appears as a featureless density in the centre of the phage capsid.

**Figure S4:**
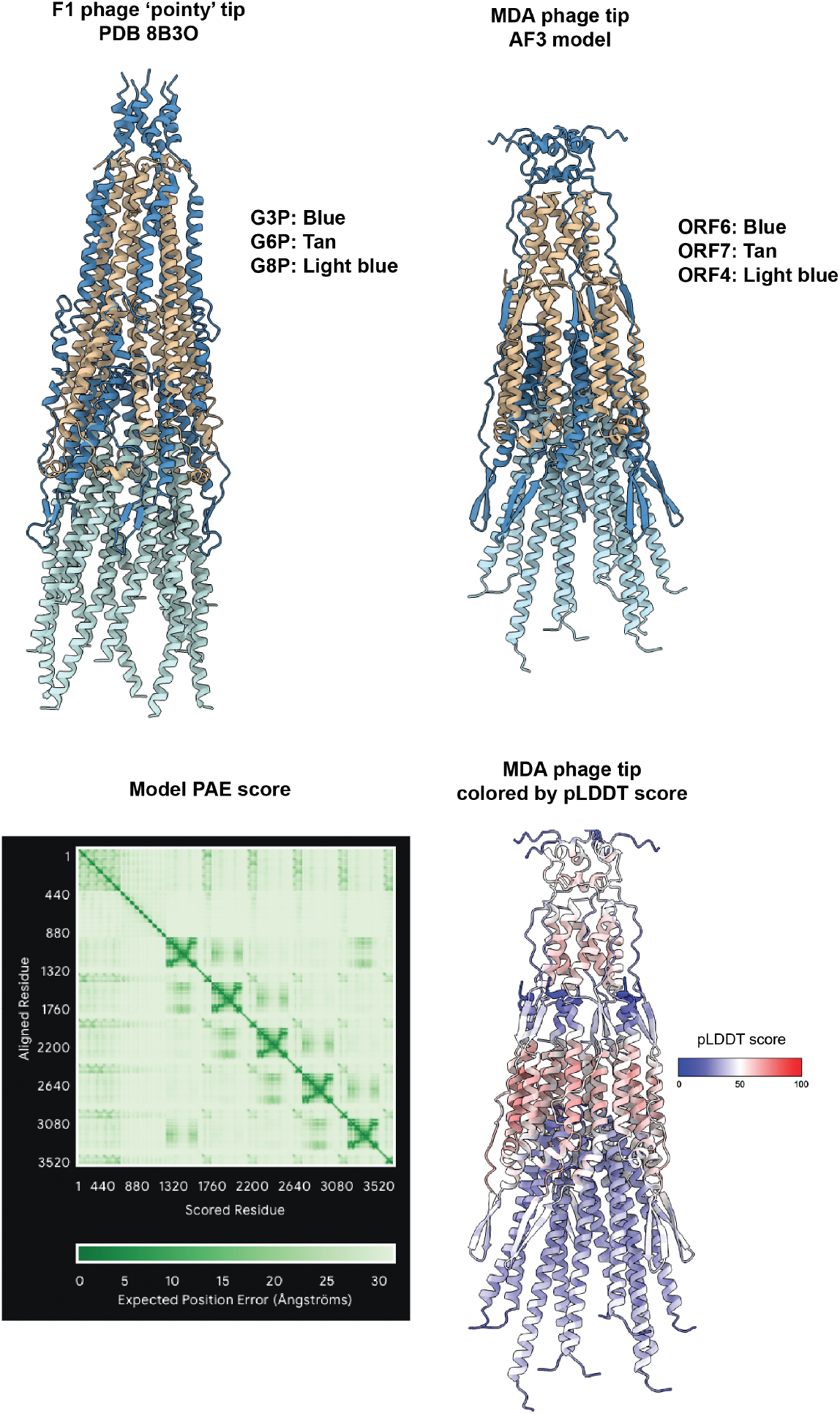
AlphaFold3 modelling of the MDA ‘pointy’ phage tip, and comparison with a cryo-EM structure of the tip of the f1 bacteriophage. This prediction containing 10 copies of the major capsid protein ORF4, 5 copies of ORF6 and 5 copies of ORF7 is architecturally highly similar to a previously solved cryo-EM structure of the f1 filamentous phage tip (PDB 8B3O (21)). N-terminal residues 1-399 of ORF6 were excluded to enable a better structural comparison. The prediction’s pTM score is 0.3 and TM score is 0.33. This proposed model will require experimental validation.

**Figure S5:**
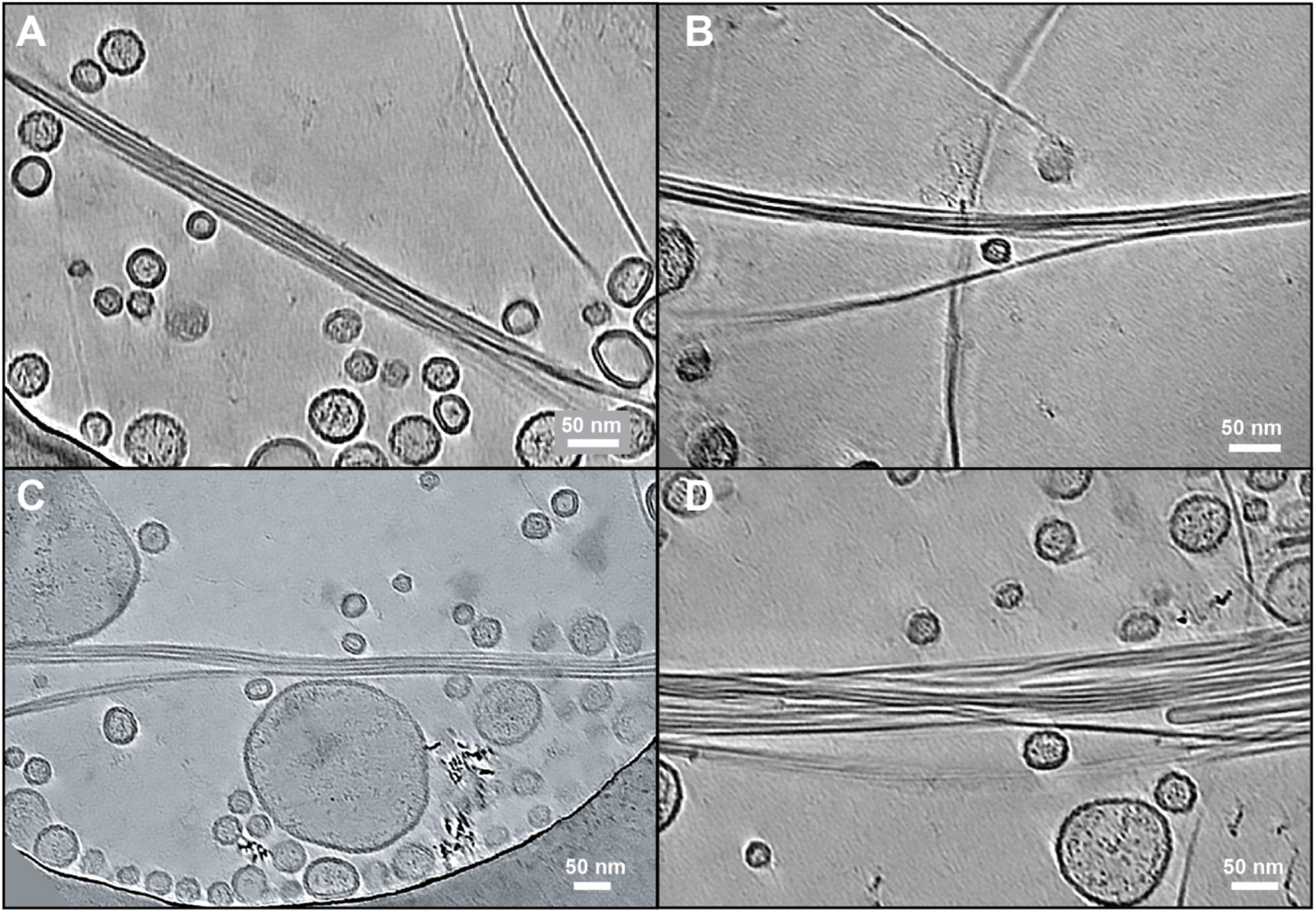
Cryo-ET of MDAΦ bundles. **A-D)** Gallery of MDA phage bundles shown as orthogonal slices from denoised tomograms. Vesicles co-purified with MDAΦ can be seen alongside phage bundles.

## Supplementary Tables

**Table S1:** Cryo-EM data acquisition and processing statistics.

|  |  |
| --- | --- |
| Data collection and processing | MDAΦ [EMDB XXXXX, PDB XXXX] |
| Microscope | Krios Titan G2 |
| Magnification | 96,000 |
| Voltage (kV) | 300 |
| Electron exposure (e<sup>-</sup>/Å<sup>2</sup>) | 41.06 |
| Defocus range (μm) | -1 to -2.5 |
| Pixel size (Å) | 0.824 |
| Symmetry imposed | C5, helical |
| Initial particle images (no.) | 260,007 |
| Final particle images (no.) | 27,134 |
| Map resolution (Å) | 3.7 |
| FSC threshold | 0.143 |
| Map resolution range (Å) | 3.5-5.3 |
| Model Refinement |  |
| Initial model used | AlphaFold2 |
| Map sharpening *B* factor (Å<sup>2</sup>) | -26.25 |
| Model composition |  |
| Non-hydrogen atoms | 1815 |
| Protein residues | 255 |
| Ligands | 0 |
| R.m.s. deviations |  |
| Bond lengths (Å) | 0.0087 |
| Bond angles (°) | 1.79 |
| Validation |  |
| MolProbity score | 2.01 |
| Clashscore | 7.20 |
| Poor rotamers (%) | 2.78 |
| Ramachandran plot |  |
| Favored (%) | 95.92 |
| Allowed (%) | 4.08 |
| Disallowed (%) | 0 |

**Table S2:** Genome type (linear versus circular) and number of lysine and arginine residues in the C-termini of the MCPs in reference genomes (18). We propose that the number of C-terminal arginine (R) or lysine (K) residues correlate with genome type, with 3-4 basic residues correlating with a circular genome and 2 basic residues correlating with a linear genome. Reference genomes to which this rule applies are marked in green, reference genomes where it does not apply are marked in tan colour.

| Genome | Protein | Genome | Number of R/K |
| --- | --- | --- | --- |
| LC035386 | LC035386.BAS49600.1 | circ | 4 |
| LC210520 | LC210520.BAX02602.1 | circ | 4 |
| NC_001418 | NC_001418.NP_040652.1 | circ | 2 |
| NC_003460 | NC_003460.NP_604425.1 | circ | 4 |
| NC_004306 | NC_004306.NP_695190.1 | circ | 4 |
| NC_004736 | NC_004736.NP_835475.1 | circ | 4 |
| NC_012757 | NC_012757.YP_002925191.1 | circ | 4 |
| AB931172 | AB931172.BAP81896.1 | circ | 3 |
| AY324828 | AY324828.AAQ94685.1 | linear | 2 |
| CP017895 | CP017895.ARP06761.1 | circ | 4 |
| GQ153916 | GQ153916.ACY07131.1 | circ | 4 |
| GQ153919 | GQ153919.ACY07161.1 | circ | 4 |
| LC066596 | LC066596.BAS04448.1 | circ | 3 |
| NC_001331 | NC_001331.NP_039603.1 | linear | 2 |
| NC_001332 | NC_001332.NP_039620.1 | circ | 4 |
| NC_001396 | NC_001396.NP_536674.1 | circ | 3 |
| NC_001954 | NC_001954.NP_047355.1 | circ | 4 |
| NC_002014 | NC_002014.NP_040575.1 | circ | 4 |
| NC_003287 | NC_003287.NP_510890.1 | circ | 4 |
| NC_003327 | NC_003327.NP_752644.1 | circ | 3 |
| NC_005948 | NC_005948.YP_031682.1 | circ | 4 |
| NC_008575 | NC_008575.YP_863924.1 | circ | 3 |
| NC_015297 | NC_015297.YP_004327583.1 | circ | 3 |
| NC_021562 | NC_021562.YP_008130278.1 | circ | 4 |
| NC_021866 | NC_021866.YP_008320409.1 | circ | 3 |
| NC_023586 | NC_023586.YP_009008129.1 | circ | 4 |
| NC_025824 | NC_025824.YP_009111302.1 | circ | 4 |
| AB572858 | AB572858.BAJ12070.1 | circ | 4 |
| AF452449 | AF452449.AAL40839.1 | circ | 4 |

## Movie captions

**Movie S1: Cryo-EM structure of the MDAΦ capsid**.

Cryo-EM density and ribbon depiction of the atomic model of the MDAΦ capsid are shown.

**Movie S2: Electron cryotomography of MDAΦ bundles**.

Shown are sequential Z-slices of a tomogram of an MDAΦ bundle.

## Notes

### Competing Interest Statement

The authors have declared no competing interest.

